# Transferrin Receptor Targeting Chimeras (TransTACs) for Membrane Protein Degradation

**DOI:** 10.1101/2023.08.10.552782

**Authors:** Dingpeng Zhang, Jhoely Duque-Jimenez, Garyk Brixi, Francesco Facchinetti, Kaitlin Rhee, William W. Feng, Pasi A. Jänne, Xin Zhou

**Affiliations:** Department of Cancer Biology, Dana-Farber Cancer Institute, Boston, MA, USA; Department of Biological Chemistry and Molecular Pharmacology, Harvard Medical School, Boston, MA, USA; Harvard University, Boston, MA, USA; Department of Medical Oncology, Dana-Farber Cancer Institute, Boston, MA, USA; Department of Medicine, Harvard Medical School, Boston, MA, USA; Lowe Center for Thoracic Oncology, Dana-Farber Cancer Institute, Boston, MA, USA; Belfer Center for Applied Cancer Science, Dana-Farber Cancer Institute, Boston, MA, USA

## Abstract

Cancer cells require high levels of iron for rapid proliferation, leading to a significant upregulation of the iron carrier protein Transferrin Receptor (TfR) on their cell surface. Leveraging this phenomenon and the exceptionally fast endocytosis rate of TfR, we introduce Transferrin Receptor TArgeting Chimeras (TransTAC), a novel molecular archetype for membrane protein degradation in cancers and other cell types. TransTACs repurpose the naturally recycling receptor TfR1 for protein degradation. To accomplish this, we utilized a combination of protein engineering strategies to redirect the target protein from recycling-endosome trafficking to lysosomal degradation. We show that TransTACs can highly efficiently degrade a diverse range of single-pass, multi-pass, native, or synthetic membrane proteins, establishing new possibilities for targeted cancer therapy.

## Introduction

Targeted protein degradation (TPD) is a rapidly growing field in drug discovery and pharmacology. Complementing traditional drug modalities, TPD molecules offer a novel therapeutic mechanism to tackle challenging targets or increase the therapeutic potential of currently used drugs^1^. While most efforts in this field have focused on small molecules for intracellular targets, inducing targeted degradation of membrane proteins has recently emerged as a new therapeutic opportunity^2^. Membrane proteins are central to a myriad of cellular functions and serve as targets for over half of all drugs^3^. Therefore, developing universal strategies to degrade membrane proteins is of exceptional interest for both basic research and therapeutic intervention purposes.

Over the last few years, several proof-of-concept strategies for membrane receptor degradation have been described. These strategies use heterobifunctional biologics that recruit a specific “effector” protein such as a membrane E3 ligase^4,5^ or a lysosome shuttling receptor^6-8^ to the protein of interest (POI) to induce lysosome-mediated protein degradation. However, the effectiveness of these biological effectors is often limited by their tissue-specific expression patterns. GalNAc-LYTAC, for instance, targets the hepatocyte-specific receptor asialoglycoprotein receptor (ASGPR), making it only suitable for treating liver disease or clearing circulating targets^6,9^, while RNF43- or ZNRF3-based methods are more effective for treating Wnt-signaling upregulated disorders where RNF43 and ZNRF3 are expressed at high levels^4,5^. Current technologies are therefore not able to cover the full spectrum of diseases, and developing alternative effectors overexpressed in different diseases and tissues would greatly expand the range of cell surface targets that can be regulated and also increase the targeting specificity.

Iron is an essential element for cells, and its transportation is facilitated by the transferrin receptor 1 (TfR1)^10,11^. Rapidly dividing cells, such as cancer cells and activated immune cells, exhibit substantially increased TfR1 expression compared to non- or slow-dividing cells due to their high demand for iron^12,13^. As a result, TfR1 is an attractive target for modulating these cell types. Several studies have investigated TfR1 as a therapeutic target for cancer with promising outcomes^12,14^, as well as for tumor imaging^15^ and targeted drug delivery^16^.

Apart from its overexpression in proliferating cells, the rapid internalization rate is another intriguing feature of TfR1. As a classical recycling receptor, extensive research has been conducted on the intracellular trafficking pathways and internalization kinetics of TfR1. TfR1 exhibits an average internalization rate of 500 molecules per cell per second, making it one of the fastest internalizing receptors known^17,18^. This exceptional speed suggests that TfR1 has the potential to serve as a carrier “effector” for inducing targeted internalization of membrane proteins.

In this study, we leveraged these two crucial features of TfR1 and employed protein engineering strategies to develop a novel technology for degrading membrane proteins. We call this technology Transferrin Receptor TArgeting Chimeras (TransTAC). TransTACs are heterobispecific antibodies that bring the POI and TfR in close proximity at the cell surface and separate them in the endosomes for lysosome-mediated degradation (**Fig. 1a**). We show that TransTACs are highly effective in degrading various types of membrane proteins, including single-pass, multi-pass, native, and synthetic receptors, showing a degradation efficiency of over 80% in various cellular systems. An intriguing characteristic of TransTACs is the fast kinetics of targeted internalization, which occurred on the timescale of minutes in the context of a chimeric antigen receptor (CAR). Moreover, TransTAC molecules are fully recombinant, versatile, and modular. These unique properties make TransTACs a promising tool for the precise manipulation of cell surface targets, particularly in cancer and immune cells, offering new opportunities for precise targeting of the extracellular proteome in research and medicine.

**Figure 1.**
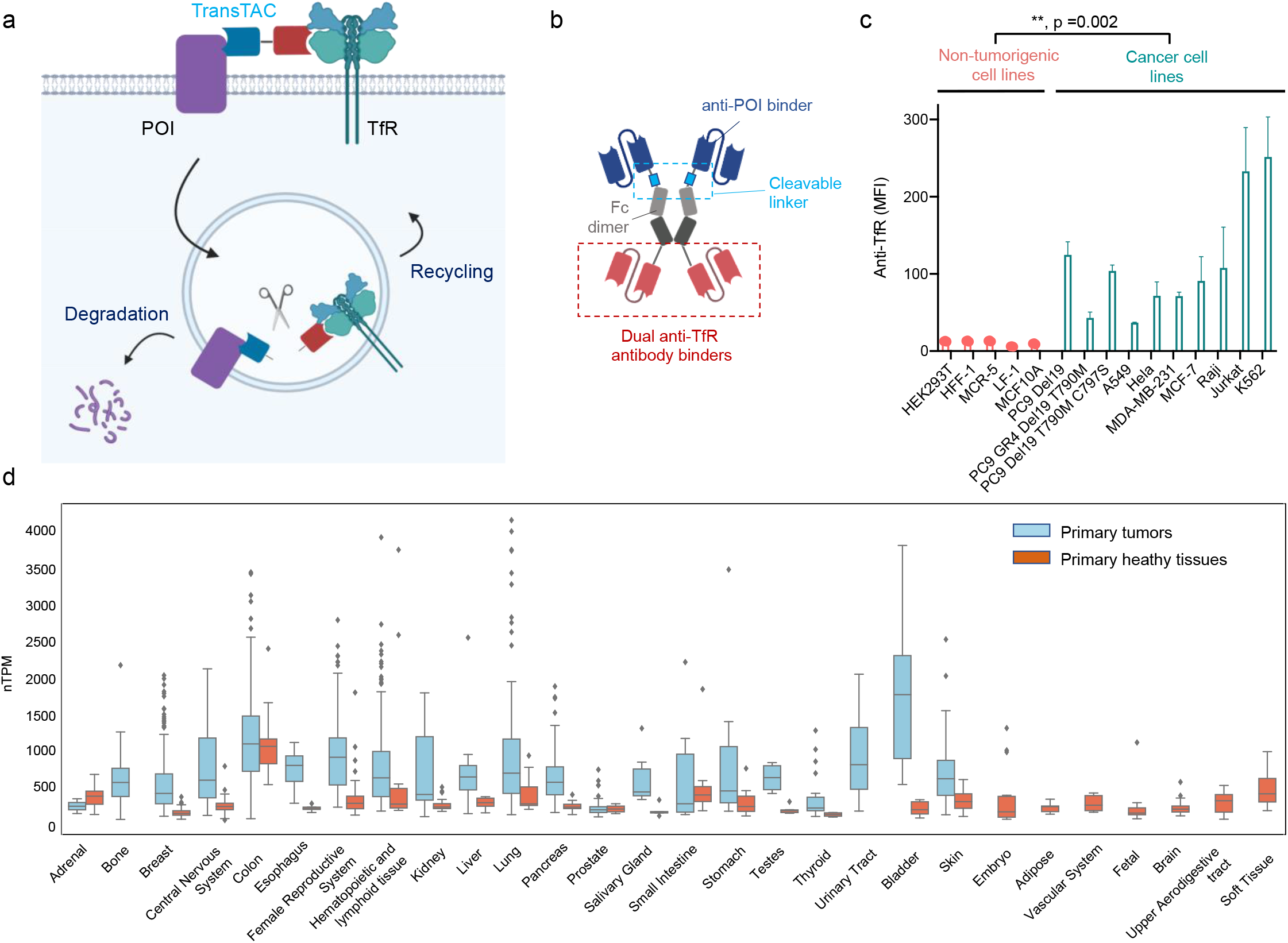
Overview of the TransTAC technology and Transferrin Receptor 1 (TfR1) expression analysis. **(a)** Schematic of TransTACs. TransTAC induces close proximity of TfR and a protein of interest (POI) at the cell surface, leading to co-internalization of the complex to the early endosomes, where a cathepsin enzyme cleaves TransTAC and separates the POI from the TfR. The POI then traffics to late endosomes/lysosomes for degradation, while TfR is recycled back to the cell surface. **(b)** Illustration of an example TransTAC protein. Key designs to make TransTACs efficient degraders include: (1) containing two anti-TfR1 binders for binding and priming a TfR1 dimer for endocytosis, (2) having a cathepsin-sensitive linker between the anti-POI binder and the Fc for endosomal cleavage to separate the POI from the recycling TfR1s, and (3) using an antibody binder instead of a native TfR1 ligand to reduce trafficking to the recycling endosomes. **(c)** Relative cell surface TfR1 expression levels across various non-tumorigenic and cancer cell lines characterized by flow cytometry. Cancer cell lines express significantly higher levels of TfR1 compared to non-tumorigenic cell lines. Data are representative of 3 independent experiments. **(d)** Relative *TFR1C* RNA expression levels in primary tumor compared to normal tissues based on the MERAV database. *TFR1C* expression is significantly higher in most tumors than the corresponding normal tissues. T-test in **Extended Data Fig. 1a** shows significance for comparing tumors to healthy tissue overall (p= 3.98e-89), and for 14 out of 19 of the individual tumor/healthy tissue pairs. Female reproductive tissues are the endometrium, cervix, fallopian tubes, myometrium, ovary, placenta, and uterus. Central nervous system (CNS) tissues are the basal ganglia, brainstem, cerebral cortex, hippocampus, spinal cord, and vestibular nuclei superior. Brain tissues are the hypothalamus, pituitary gland, thalamus, ganglia, and ganglion nodose. Full sample IDs and labels are available in **SI appendix table**. Error bars represent standard deviations. Fig. 1a, b are created with BioRender.com.

### TfR1 expression is upregulated in cancer cell lines, primary tumors, and activated T cells

Our rationale for developing the TransTAC technology is that overexpression of TfR1 in malignant tissues may enable enhanced tumor targeting specificities of the degrader. Thus, we sought to validate cancer-specific TfR1 overexpression and analyze its tissue distribution. Cell surface TfR1 levels were measured on ten cancer cell lines and five non-tumorigenic cell lines, including lung, breast, cervical, lymphoma, and leukemia cancers (**Fig. 1c**). TfR1 overexpression was observed on all cancer cell lines compared to normal cell lines with statistical significance (p = 0.002). Among different cell types, Jurkat and K562 leukemia cell lines exhibited the highest TfR1 expression, with more than 20-fold greater TfR1 expression compared to normal cells. Similarly, A549 (EGFR wildtype) and PC9 (EGFR mutant) lung cancer cell lines showed 3 to 11-fold greater TfR1 expression than normal cells.

We further showed TfR1 expression is upregulated in primary tumors compared to primary healthy tissues, by performing a transcriptomics analysis of *TFR1C*, the gene for TfR1 (**Fig. 1d, Extended Data Fig. 1a**)^19^. Microarray transcriptomics data of *TFR1C* was obtained from the MERAV database^20^. Paired tissue analysis was performed using a custom python script (**SI Appendix**). *TFR1C* expression is statistically significantly increased in cancers overall (p = 3.98e-89) and in 14 out of 19 specific tissues, including breast, lung, pancreas, liver, bladder, skin, esophagus, thyroid, testes, stomach, salivary gland, kidney, central nerves system (CNS), and female reproductive system tumors. Adrenal samples exhibited higher *TFR1C* levels in healthy tissues, but without statistical significance. Small intestine, prostate, colon, and hematopoietic and lymphoid samples had higher overall expression of *TFR1C* in tumor than normal tissues, but without statistical significance. The lack of statistical significance for hematopoietic and lymphoid samples may be due to heterogeneity of various subsets of cell types. Among all measured healthy tissues, colon tissues had the highest basal *TFR1C* expression.

To investigate whether TfR1 could also be a potential target for immune cell modulation, the DICE dataset was analyzed, which contains gene expression profiles of human immune cells isolated from blood samples of healthy donors^21^. While most immune cells express a low level of TfR1s, an approximately 6-fold higher TfR1 expression in activated CD4 and CD8 T cells was observed compared to inactivated T cells, a level comparable to the TfR1 levels in some malignant tissues, indicating TfR1 is also a promising target for modulating activated T cells (**Extended Data Fig. 1b**,**c**). Together, our analysis of TfR1 expression in cell lines, primary cancer and healthy tissues, and primary immune cells reveal the potential of targeting TfR1 for specific modulation of tumor cells and activated T cells. Our transcriptomic analysis provides a detailed comparison of TfR1 expression in specific tissues, serving as a roadmap for future selection of disease indications for our technology and beyond.

### The Rational Design and Optimization of TransTAC Degraders

We next set out to design TransTACs. Generally, TransTACs are recombinant proteins consisting of anti-POI and anti-TfR1 binders to bring the POI and TfR1 in close proximity on the cell surface. However, we identified three crucial design principles that are important for making TransTACs efficient degraders: (1) *dimeric* TransTACs are more effective at driving protein internalization than monomeric ones, (2) a *cathepsin-sensitive linker* is necessary for POI separation from TfR1 and lysosomal trafficking, and (3) using an *antibody binder* to target TfR1, rather than a native transferrin (TF) ligand, can minimize POI trafficking to the recycling endosomes (REs) and increase degradation efficiency (**Fig. 1b**). Here, we describe how these principles were discovered, using the anti-CD19 chimeric antigen receptor (CAR)-TransTAC as an example.

Our first discovery was that a dimeric TransTAC is more effective than a monomer in internalizing CAR. Two versions of TransTAC were made: v0.1, a knob-in-hole (KIH) Fc heterobispecific, and v0.2, a homodimeric Fc fusion (**Fig. 2a**). The TfR1 ligand TF was used for binding TfR1. A stable CD19 ectodomain variant, CD19NT.1, evolved by yeast display, was used for binding CAR^22^. A CD19NT.1-Fc fusion showed a homogeneous band in the SDS-PAGE gel, whereas a native CD19 ectodomain-Fc showed aggregations, indicating that using the engineered binder was necessary (**Extended Data Fig. 2**). CAR-TransTACv0.1 and 0.2, along with a control lacking the TF ligand, were recombinantly expressed and incubated with Jurkat T lymphocyte cells expressing a Myc-tagged CAR overnight (**Fig. 2b**). Cell surface CAR levels were measured using an anti-Myc antibody. Both v0.1 and v0.2 substantially decreased cell surface CAR levels, with a maximal percent decrease (D_max_) of approximately 60% for v0.1 and 80% for v0.2 (**Fig. 2c**). CAR internalization was dependent on binding to TfR1, as the control CD19NT.1-Fc did not result in a decrease in cell surface CAR levels. Intriguingly, a hook effect was observed with v0.1, but not v0.2. This is potentially because avidity could promote cooperative binding and hence simultaneous engagement of POIs and TfR1s at the cell surface, reducing the hook effect within a broader concentration range^23^. Overall, the observation of dimeric TransTACs being more effective is consistent with the notion that TfR1 is a homodimeric receptor and requires binding of two TFs to fully prime its physiological functions^24^.

**Figure 2.**
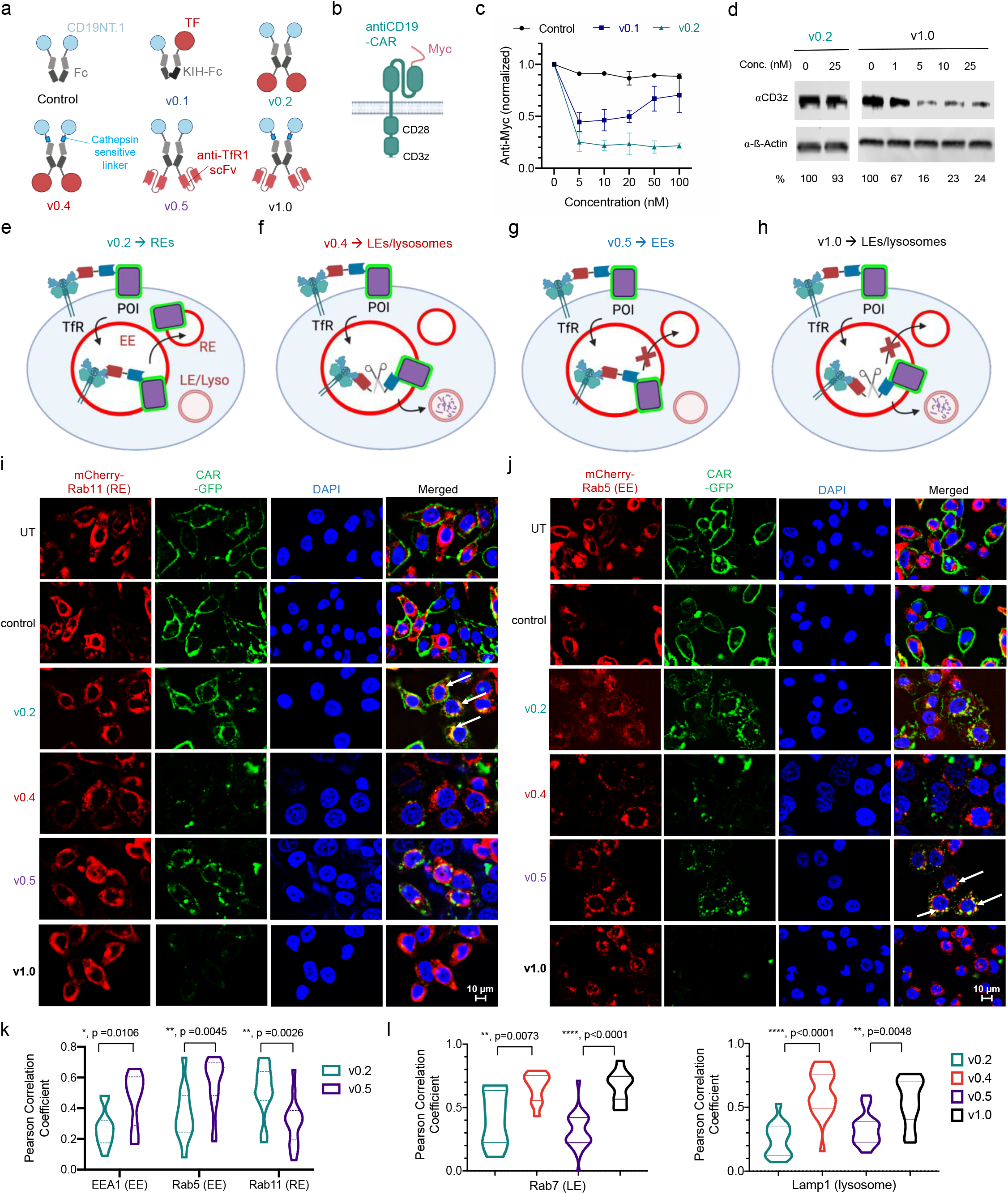
TransTAC degrader engineering. **(a)** Schematic of CAR-TransTACs and control. TransTACv0.1 consists of a single CD19NT.1 domain, a single TF, and a knob-in-hole (KIH) Fc, v0.2 consists of two CD19NT.1s, two TFs, and a homodimeric Fc that connects the binders, v0.4 contains a cathepsin-sensitive linker between CD19NT.1 and Fc, v0.5 contains an anti-TfR1 scFv for TfR1 binding, v1.0 contains both the anti-TfR1 scFv and the cathepsin-sensitive linker. **(b)** Schematic of a Myc-tagged anti-CD19 CAR receptor. **(c)** Flow cytometry measurements of cell surface CAR expression levels in CAR-Jurkats treated with TransTACv0.1, v0.2, and control. TransTACv0.2 results in higher CAR clearance from cell surface than v0.1 and no hook effect. Data are representative of 2 independent experiments. Error bars represent standard deviations. **(d)** Characterization of whole-cell CAR levels by Western blot in CAR-Jurkats treated with TransTACs. TransTACv1.0 degrades approximately 80% of CAR; v0.2 did not result in significant CAR degradation. **(e-h)** Schematics showing different TransTACs alter intracellular trafficking of the POI. Cleavage of the cathepsin-sensitive linker in v0.4 and v1.0 leads to separation of POI from TfR1, hence enhancing LE/lysosomal trafficking of the POI and degradation; the anti-TfR1 scFv in v0.5 and v1.0 reduces trafficking of the complex to the REs, hence increasing the proportion of POI in EEs and subsequent proteolytic processing, when a cleavable linker is present. **(i, j)** Representative fluorescence images of HeLa cells co-expressing CAR-GFP (green) and endosomal/lysosomal markers-mCherry (red) treated with various TransTAC molecules. Cell nucleus is stained with Hochest (blue). Untreated (UT) or control-treated cells had CAR-GFP localized at the cell membrane. v0.5 and v1.0 led to efficient degradation of CAR-GFP, manifested by the significantly lower GFP signals. v0.2-treated cells predominantly trafficked CAR to the REs, showing co-localization of CAR-GFP with mCherry-Rab11 (white arrows). v0.5-treated cells trafficked CAR to the EEs, showing co-localization of CAR-GFP with mCherry-Rab5 (white arrows). **(k)** Pearson correlation analysis of CAR-GFP colocalization with the Rab5 (EE), EEA1 (EE), and Rab11 (RE) markers. T-tests show Rab5, EEA1, Rab11 colocalization with CAR are statistically different for cells treated with v0.2 vs. v0.5. **(l)** Pearson correlation analysis of CAR-GFP colocalization with the Rab7 (LE) and Lamp1 (lysosome) markers. T-tests show Rab7 and Lamp1 colocalization with CAR are statistically significant for v0.2 vs. v0.4, and v0.5 vs. v1.0. For k and l, the number of cells used for each analysis are as follows: For v0.2, N=12, N=12, and N=13 for the EEA1, Rab5 and Rab11 markers, respectively. For v0.5, N=10, N=22, and N=15 for the EEA1, Rab5 and Rab11 markers respectively. For v0.2, v0.4, v0.5, and v1.0 with the Lamp1 marker N=16, N=21, N=13, and N=13, respectively. Fig. 2a, b, e-h are created with BioRender.com.

Interestingly, despite the effective cell surface removal of CAR, TransTACv0.2 did not result in CAR degradation, as shown by whole-cell lysate western blots (**Fig. 2d, Extended Data Fig. 3a, b)**. To understand the subcellular destination of the internalized CAR, we stably expressed CAR-GFP and mCherry tagged to different endosomal and lysosomal markers, including Rab5+ or EEA+ for early endosomes (EEs), Rab7+ for late endosomes (LEs), Rab11+ for REs, and Lamp1+ for lysosomes, in a HeLa cell line (**Fig. 2i, j, Extended Data Fig. 4)**^25^. Fluorescence microscopy imaging of v0.2-treated cells showed co-localization of CAR-GFP with Rab11, indicating that the internalized CAR trafficked to the REs with v0.2 (**Fig. 2i, white arrows**).

To promote CAR entry into the degradative pathway and prevent sorting into the recycling pathway with TfR1, we incorporated a cathepsin-sensitive linker for proteolytic processing in the EEs^26^. We reasoned that to prevent CAR from being sorted into the recycling pathway and instead promote its entry into the degradative pathway, it would be necessary for CAR to disengage from TfR1 in the EEs (**Fig. 2e,f**). A cathepsin-sensitive sequence was inserted in TransTACs, either between Fc and TF (v0.3), or between CD19NT.1 and Fc (v0.4) **(Fig. 2a, Extended Data Fig. 3a)**. Western blot analysis of CAR-Jurkat cells treated with v0.4 showed approximately 40-50% degradation of CAR (**Extended Data Fig. 3c**). Furthermore, HeLa cells expressing CAR-GFP treated with v0.4 showed a substantial reduction in GFP fluorescence (**Fig. 2i, j, Extended Data Fig. 4a-c**), showing the new linker design enabled protein degradation.

To determine whether trafficking to LEs and lysosomes was enhanced, we collected fluorescence images for Pearson correlation analysis of GFP and mCherry. A significant increase was observed in CAR-GFP/Rab7 and Lamp1 co-localization, markers for LEs and lysosomes, compared to co-localization with Rab11, validating our hypothesis that endosomal sorting to the degradative pathway was enhanced with the linker variation (**Fig. 2i, Extended Data Fig. 4d, 5**). Interestingly, the degradation efficiency was substantially lower with v0.3 (**Extended Data Fig. 3c**). This is possibly because CD19NT.1 remains linked to the Fc after cleavage of TransTACv0.3, and Fc can subsequently mediate protein recycling via the FcRn pathway^27^.

We further screened a panel of linkers to identify optimal protease substrates in EEs^28^. Fourteen linkers containing single or combined cathepsin cleavage motifs were incorporated into the CAR-TransTAC and characterized using western assays (**Extended Data Fig. 3d**,**e**)^29^. Two of the best linkers from this panel were selected to develop new TransTACs.

Lastly, we substituted TF with an anti-TfR1 single-chain Fv (scFv) identified from phage display. Our rationale was that TF may contain specific structural or signaling elements that promote trafficking of internalized proteins to the REs. Therefore, using a synthetic antibody may reduce trafficking of the internalized complex to the REs, resulting in an increased percentage of POIs that stay in the EE compartments for proteolytic processing and trafficking to the degradative pathway (**Fig. 2g, h**). Two new TransTACs were generated: v0.5 containing the anti-TfR1 scFv binder but no cleavable linker, and v1.0 containing both the anti-TfR1 scFv and the cleavable linker (**Fig. 2a**). Remarkably, cells treated with v0.5 showed strong colocolization of CAR-GFP with EE markers Rab5 and EEA, but not RE marker Rab11 (**Fig. 2i, j, Extended Data Fig. 4a)**. Pearson colocalization coefficients of the corresponding markers were statistically different from cells treated with TransTACv0.2, which contains the TF ligands (**Fig. 2k, Extended Data Fig. 4d)**. Additionally, protein degradation efficiency was substantially increased. With v1.0, over 80% CAR degradation was observed in the Western, and nearly no CAR-GFP signals were observed in fluorescence images (**Fig. 2d, i, j, Extended Data Fig. 4a-c)**. Additionally, the anti-TfR1 scFv substitution also increased the yield of the protein by approximately seven-fold, making the expression level of TransTACs similar to that of conventional antibodies.

Taken together, our rational protein engineering efforts have successfully rewired the intracellular trafficking of the target protein in the endosomal-lysosomal pathway, resulting in a novel protein degrader design, TransTACv1.0, that leads to >80% CAR degradation. Notably, several earlier versions of CAR-TransTACs have the ability to effectively eliminate CAR from the cell surface through an endosomal trapping mechanism, offering versatile possibilities of membrane protein regulation (**Fig. 2a**).

CAR-T cell therapy has demonstrated significant potential in treating hematologic malignancies. However, its broad application is limited by the risk of potentially fatal side effects, such as cytokine release syndrome (CRS), caused by the overactivation of CAR-T cells^30,31^. To this end, CAR-TransTACs can potentially serve as a modular protein OFF switch to fine tune CAR-T cell activity and to manage its associated toxicities (**Extended Data Figure 6a**). As a proof of concept, we showed that CAR-TransTACv0.4 can effectively inhibit human primary CAR-T cell IFN-γ secretion, giving an impressively low IC50 of 0.4 nM (**Extended Data Figure 6b, c**). Additionally, it reversibly regulated CAR-T cell-mediated tumor killing activities (**Extended Data Figure 6b, d, e**). To the best of our knowledge, CAR-TransTAC is the first recombinant protein-based OFF-switch for CAR-T cell regulation that does not need additional modifications to the CAR-T cells. Unlike splitCAR- or other circuit rewiring-based methods^31^, CAR-TransTACs do not require genetic engineering and are, therefore, readily adaptable to a variety of CAR-T cell therapies, both approved and in development.

### Expansion of TransTAC-addressable targets

We next established the generalizability of TransTAC degraders (**Fig. 3a**). We aimed to include targets that have diverse structures and functions present on different cell types, such as native vs. synthetic, single-pass vs. multi-pass transmembrane, and cancer vs. immune cell targets.

**Figure 3.**
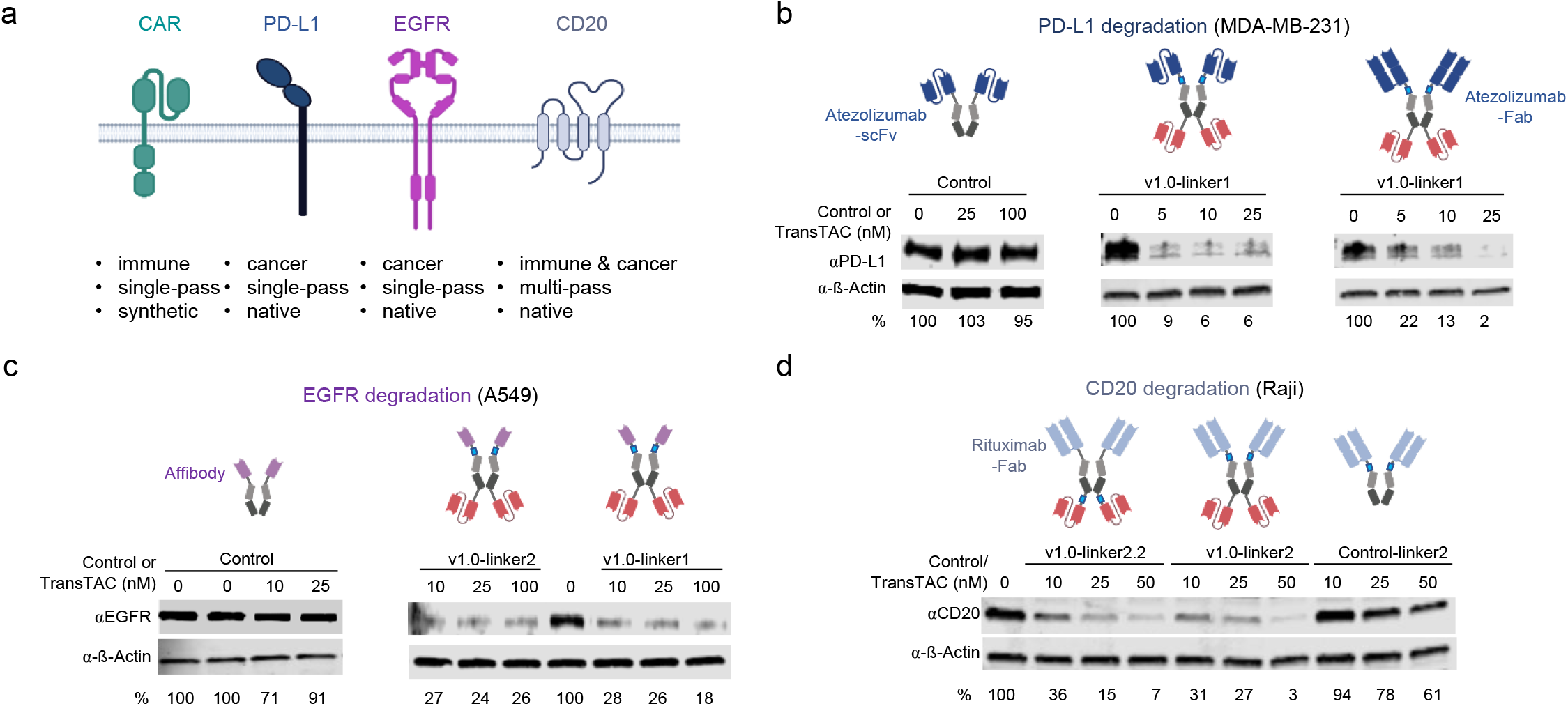
Developing TransTACs degraders for various proteins. **(a)** Schematic of membrane proteins targeted by TransTACs in the present study. These targets are either synthetic, or native, single- or multi-pass proteins expressed on the cancer or immune cell surface. **(b)** PD-L1 degradation by TransTACs in MDA-MB-231 breast cancer cells analyzed by Western blot. A scFv or Fab format of atezolizumab is used as the PD-L1 binding moiety. **(c)** EGFR degradation by TransTAC in A549 lung carcinoma cells. An affibody is used as the EGFR binding moiety. **(d)** CD20 degradation by TransTAC. A Fab format of rituximab is used as the CD20 binding moiety. Fig. 3a and illustrations of TransTACs in 3b-d are created with BioRender.com.

Our first target is programmed death-ligand 1 (PD-L1), an immune checkpoint receptor ligand, downregulation of which can enhance anti-tumor T cell activity^32^. Despite having improved clinical outcomes across tumors, many patients still do not benefit from monoclonal antibodies targeting the PD-1/PD-L1 axis, making new mechanisms for targeting PD-L1 highly desirable. A PD-L1-TransTAC was created using a fragment antigen binding (Fab) or single-chain variable fragment (scFv) of atezolizumab as the PD-L1 binding domain (**Fig. 3b**) ^33^. Up to 98% PD-L1 degradation was observed in MDA-MB-231 breast cancer cells treated with PD-L1-TransTACs, while control groups lacking the anti-TfR1 scFv binder or containing TransTACv0.2 and v0.4 with a TF ligand showed no or little PD-L1 degradation (**Fig. 3b, Extended Data Fig. 7a)**.

Next, we aimed to target EGFR, a receptor tyrosine kinase that plays a critical role in the development and progression of various types of cancers such as lung and brain^34^. An EGFR-TransTAC was created with an affibody as the EGFR binding domain (**Fig. 3c**)^35^. A549 lung carcinoma cells treated with EGFR-TransTACv1.0s showed up to 80-90% reduction of EGFR, whereas control groups exhibited little to no degradation (**Fig. 3c, Extended Data Fig. 7b**). Different linkers in v1.0 resulted in varying degrees of EGFR degradation, while v0.2 had no effect (**Extended Data Fig. 7b**). These results were consistent with the observations with the CAR-TransTAC variants, validating the importance of those modifications made to improve TransTAC efficiency.

Cluster of differentiate 20 (CD20) is a B cell-specific surface marker with four transmembrane domains and an unknown function^36^. Knocking down cell surface CD20s with a degrader could be valuable for functional studies of CD20. Using a scFv and a Fab format of rituximab, the first clinically approved antibody directed against CD20 in the setting of non-Hodgkin lymphoma, we created a CD20-TransTAC (**Fig. 3d**)^37^. Treatment of Raji cells, a human B lymphoblastoid cell line, with the CD20-TransTAC resulted in up to 97% reduction of CD20, while control groups led to no or significantly less degradation (**Fig. 3d, Extended Data Fig. 7c**).

Together, the successful generation of degraders against all four targets demonstrates the modularity and generality of the TransTAC design. Importantly, high potencies were observed for all four targets, highlighting the efficiency of targeted degradation using TransTACs.

### Kinetics, structure-activity relationship, mechanism, and *in vivo* characterization of TransTACs

We then studied the kinetics of TransTAC-mediated protein internalization. A time-course measurement of cell surface CAR levels in response to TransTACs was performed (**Fig. 4a**). A rapid elimination of CAR from the cell surface was observed, with only 17% remaining after 10 minutes and 13% after 20 minutes of treatment with TransTACv1.0. Furthermore, this response was long-lasting, with 10% of CAR observed at the cell surface after 3 hours with v1.0. This fast and sustained protein downregulation highlights TransTACs as a promising research tool for knocking down cell surface proteins as an alternative to genetic methods.

**Figure 4.**
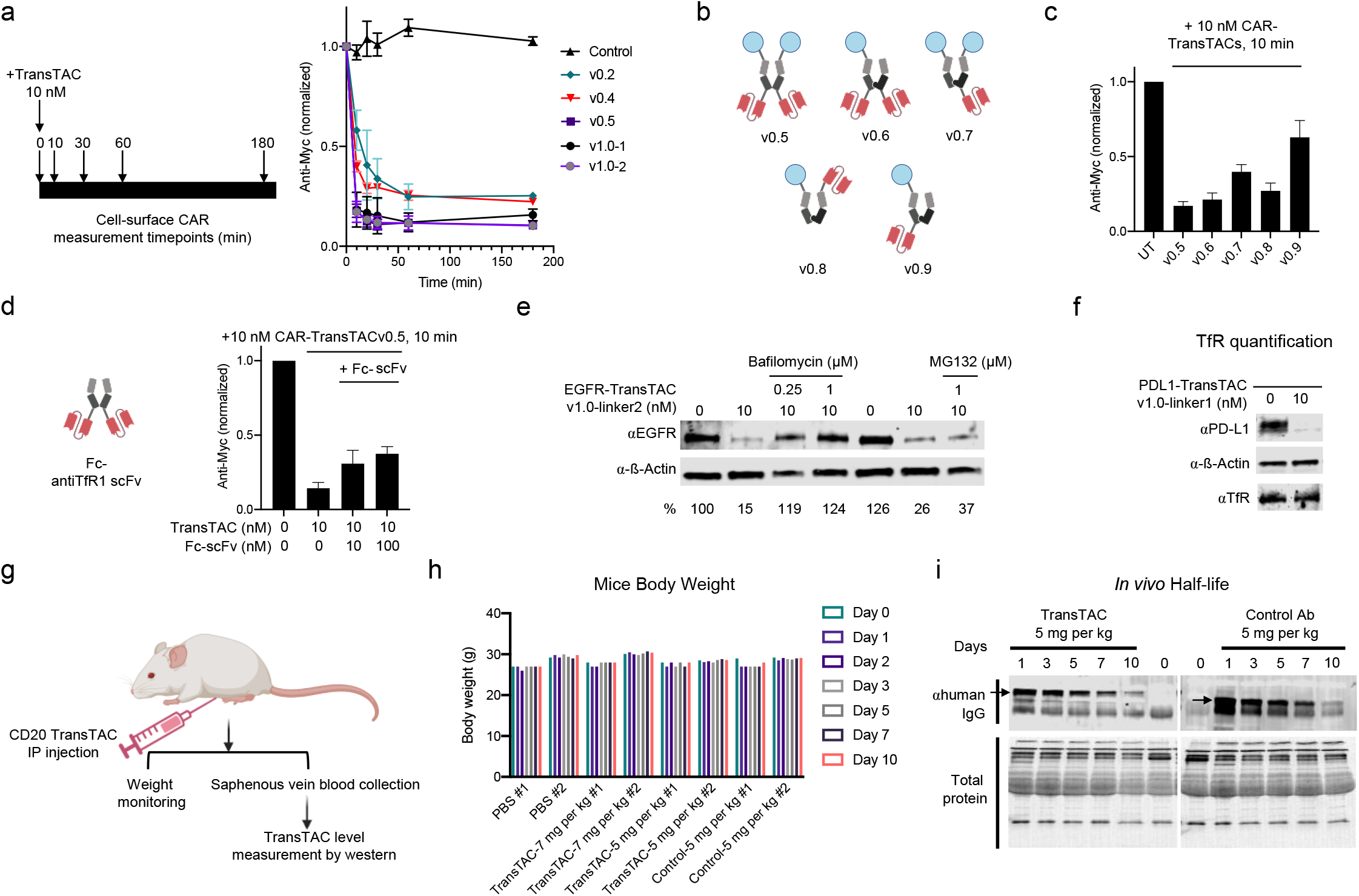
Structure-activity relationship of TransTACs, mechanisms, and *in vivo* characterizations. **(a)** Time-course measurement of cell surface CAR levels in CAR-Jurkats treated with TransTACs, revealing the fast kinetics of TransTAC-mediated CAR internalization. **(b)** Schematics of CAR-TransTAC variants consisting of one or two copies of anti-POI and anti-TfR1 binders in different protein geometries. **(c)** Cell surface CAR level measurements in CAR-Jurkats treated with CAR-TransTAC variants outlined in (b). The results highlight the impacts of having two vs. one TfR1 binders (v0.5 vs v0.7) and geometry (v0.8 vs. v0.9) in modulating protein internalization. Data are representative of 3 independent measurements. **(d)** Competition assay with a Fc-anti-TfR1 scFv fusion protein. Concentration-dependent reduction of CAR internalization is observed with Fc-anti-TfR1 scFv, proving internalization is mediated through TfR1. Data are representative of 3 independent measurements. **(e)** Study of underlying degradation pathways with TransTACs. Intact lysosomal function is critical for degradation, as degradation is fully inhibited by bafilomycin in A549 cells treated with EGFR-TransTACs. **(f)** Whole-cell TfR1 level measurement with TransTAC treatment. TfR1 level stays consistent while PD-L1 is degraded in MDA-MB-231 cells treated with PD-L1 TransTAC. **(g)** Schematic of mouse experiments to assess TransTAC safety and serum half-life via IP injection. **(h)** Weight monitoring of mice over time after TransTAC or control IgG injection. Results reveal no observable effects on mouse weight over time, showing molecules are well tolerated. N=2 per treatment group. **(i)** Western blots of plasma levels of CD20-TransTAC and control IgG over time, with quantification of data in **Extended Data Fig. 8**. Results reveal TransTAC remained in plasma up to 10 d after injection. N=2 per treatment group. Illustrations in Fig. 4b, d, g are created with BioRender.com.

To further understand how the number of binders and geometry of TransTACs influence its behavior, we generated and tested four CAR-TransTACv0.5 variants, v0.6-v0.9, each containing one or two copies of CD19NT.1 or anti-TfR1 scFv (**Fig. 4b**). Our findings revealed that having two anti-TfR1 binders was more critical than having two CD19NT.1s in enhancing CAR internalization (v0.6 vs. v0.7, **Fig. 4c**). This highlights the importance of dual binding to a dimeric TfR1, rather than having two anti-POI binders, in creating a potent TransTAC. It also indicated that TransTAC-mediated CAR internalization was not the result of CAR crosslinking. Additionally, we observed significant differences in the internalization efficiency for molecules with different geometries, indicating that the geometry of the tertiary complex structure plays a role in influencing TransTAC efficiency (v0.8 vs. v0.9, **Fig. 4c**). Furthermore, we generated an Fc-anti-TfR1 scFv molecule as a competitor for TfR1 binding and observed a dose-dependent decrease of CAR internalization in the presence of the competitor in solution with TransTACv0.5 treatment (**Fig. 4d**). This observation further validates that TransTACs function through a TfR1-dependent mechanism. These structure-activity relationship analyses offer valuable insights to guide future TransTAC designs.

We next investigated the cellular mechanism underlying TransTAC-mediated protein degradation. Two primary pathways involved in the degradation of cellular proteins were tested: the lysosomal pathway and the proteosome pathway^38^. A549 cells were treated either with bafilomycin, a vacuolar proton pump inhibitor that inhibits lysosomal acidification^39^, or MG132, a proteasomal inhibitor^40^. We observed 1 μM bafilomycin prevented TransTAC-mediated EGFR degradation, whereas 1 μM MG132 had a much less significant effect (**Fig. 4e**). These results show that intact lysosomal function was essential for TransTAC-mediated protein degradation.

To determine whether TfR1 level remains consistent or reduced with TransTAC treatment, we characterized whole-cell TfR1 expression using Western blotting assay with a PD-L1-TransTAC in the MDA-MB-231 cell line. No change in TfR1 level was observed, which is in clear contrast to the loss of PD-L1 in the same assay (**Fig. 4f**). This result validates our hypothesis that the POI was separated from TfR1 before being routed to degradation, while TfR1 is recycled.

Lastly, we asked whether TransTACs would be well tolerated and have similar antibody clearance to IgGs *in vivo*. We intraperitoneally injected 5 or 7 mg/kg (body weight) CD20-TransTAC or 5 mg/kg IgG control into nude mice (**Fig. 4g**). No significant weight changes were observed with either the TransTACs or the control (**Fig. 4h**). Western blotting analysis of plasma antibody levels revealed that the TransTAC remained in plasma up to 10 days after injection, which is comparable to the reported half-life of IgGs in mice (**Fig. 4i, Extended Data Fig. 8**)^41^. It was known that the anti-TfR1 antibody we used is cross-reactive with mouse TfR1. Together, these results demonstrate that TransTACs are well-tolerated, have favorable pharmacokinetics, and are not being rapidly cleared despite cross-reactivity with mouse cells.

## Discussion

Membrane proteins, accounting for approximately one-third of all human proteins, play pivotal roles in numerous cellular functions^3^. The ability to precisely down-regulate these proteins, ideally with temporal resolution, is crucial for studying and manipulating their physiological and disease-related functions. In this study, we introduced TransTAC, a novel molecular archetype for degrading membrane proteins. TransTACs achieved impressive degradation efficacies for various structurally and functionally diverse proteins, resulting in >80% maximal degradation across different cancer cell lines and targets. Notably, we demonstrated the first examples of using a bispecific antibody to degrade a multi-pass membrane protein, CD20, and a synthetic receptor, CAR, extending beyond targets previously explored in other studies. TransTACs are fully recombinant, easily produced, and compatible with various protein binding moieties, including Fab, scFv, affibody or protein ectodomains. These findings underscore the modularity of TransTAC designs.

A distinguishable advantage of TransTAC is its potential tumor specificity. Our analysis of TfR1 expression supports previous findings that TfR1 is significantly upregulated in cancer cells. Consequently, our developed TransTACs may enable selective targeting of cancer cells while minimizing toxicity to normal cells, addressing the off-target effects commonly associated with traditional cancer treatments.

TransTAC is the first protein degrader design to hijack a naturally recycling pathway, rather than a lysosome-shuttling ligand, for targeted protein degradation. Achieving this required a combination of protein engineering approaches, such as utilizing protein dimers, incorporating a cleavable linker, and employing an antibody instead of a natural ligand for effector binding. These engineering strategies proved to be remarkably effective, and we believe they may be generalizable to other cell surface effectors, broadening the potential scope of cell surface effectors viable for protein degradation.

The TransTAC technology is versatile and adaptable. By employing specific variants of the molecules, researchers can induce either endosomal trapping or lysosomal degradation of targets, allowing for customizable and modular manipulation of membrane proteins. Additionally, a distinguishing feature of TransTACs is their ability to rapidly control protein internalization, on a timescale of minutes in the context of CAR. This characteristic offers immense potential for fundamental research focused on understanding the temporal regulation of membrane protein functions and associated cell signaling pathways. In summary, TransTAC represents a novel protein degrader design for targeting cell surface proteins. Its tumor and immune cell specificity, genetic tractability, and rapid internalization kinetics promise to have a substantial impact on both fundamental research and therapeutic applications.

